# A Simple Subject Independent Channel Selection in EEG for Motor Imagery Task

**DOI:** 10.64898/2026.06.26.734867

**Authors:** Raghav Dev, Sandeep Kumar, Tapan Kumar Gandhi

## Abstract

Classification of motor imagery (MI) tasks through EEG is valuable in brain-computer interfacing and rehabilitation engineering. EEG channels selection for MI task classification is well discussed problem and is challenging due to its combinatorial nature. Most of the existing methods are subject and task-dependent. This paper introduces a subject-independent EEG channel selection. The proposed approach consists of two stages. First, we rank channels based on their divergence from a reference channel Cz. We hypothesize that channels less divergent from Cz are more relevant for MI task classification. In the second stage, we employ a three-stage feature selection and classification model to evaluate the selected channels. It consists of a bandpass filter, followed by common spatial pattern (CSP) filter and three classifiers viz. SVM, 1-NN and 5-NN. Two publicly available datasets viz. PhysioNet and BCI Competition III IVa datasets have been used to assess the method. It performs 15.21% more than 3Cs and just 2.91% less than all-channels accuracy with as few as 20/118 channels on BCI Competition data and 19.64% more than 3Cs on the PhysioNet dataset with 16/64 channels. Empirical comparison implies that the method performs better than classical models such as CSP Rank, fishers rank, and normalized mutual information, significantly. Results support that our hypothesis that divergence between channels and a reference channel Cz can be used as a ranking measure for channel selection.

## I. Introduction

**E**lectroencephalography (EEG) is a cost-effective, easy to use, and rich with temporal dynamics modality to study the human brain [1]. EEG channel selection for various cases such as motor imagery (MI) tasks, epileptic seizure detection, source localization, and emotion recognition is a well-researched topic and crucial for brain-computer interface applications [2]–[6]. We believe, there is scope for improvement on at least two fronts in existing channel selection models, and in this work, we have made an attempt to hammer them both up to some extent. The first issue is that most of the channel selection models are subject and task dependent and therefore need to be reconfigured and sometimes reinvented to be suited for subject-to-subject and task-to-task use. This makes these models impractical up to a certain extent and therefore need a principled approach to address it. The second issue we find interesting on this problem is that the literature is split on the question of whether a smaller number of channels outperforms all the channels on account of classification accuracy.

Channel selection is essentially an NP-hard problem since it is combinatorial [7] and it is difficult to find and assess the optimality of the channel selection methods [8]. Therefore we need priors and assumptions to narrow down the search. One common assumption in channel selection is that if a channel contributes to increasing classification accuracy more than another, it is better and more relevant [9]–[12]. However, there is no theoretical reason to believe that just because a channel doesn’t contribute to classification accuracy, it is redundant or noisy. It might be useful for generalizability. Contrary to this, the inclusion of noise in signals has the potential to enhance the accuracy of classification [13]. Another commonly made assumption is that an effective set of electrodes should be in close proximity to certain crucial electrodes [14], [15]. However, Helmholtz theorem [16] and analysis on electrode sensitivity [17], [18] based on it, state otherwise because of the shunting effect. Therefore, we think, the priors and assumptions to narrow down the search are at the heart of this problem and must be based on established notions.

Various other (than the distance between the electrodes) distance-based measures have been widely used for channel selection [19]. Examples such as normalized mutual information (NMI) among channels [20] and mutual information between channels and labels [21]–[24] have been used as the distance parameter to rank and select channels for the classification of the tasks. However, selected channels using mutual information based models are dependent on labels and therefore lack generalizability. Similarly, in [20], researchers used NMI to find the divergence between the channels, however, their channel selection algorithm is highly dependent on tasks. The other issue with existing NMI based models [20] is that it has been evaluated between every couple of channels. But if two channels are noisy, the NMI between those will be high too, and hence NMI between all couple of channels is not the best way to deal with channel selection. Similar is the issue with correlation based channel selection proposed in [25]. In this work, we have attempted to address it.

Classical methods such as recursive feature elimination [26] are largely subject and task specific, since, the channel selection depends on the classifier’s loss function (margin of support vector machines (SVM)). Similarly, spatial filters based channel selection models [27]–[31] are task dependent since filters have to be derived for each label. Recent works such as Riemannian channel selection models [32], [33] are subject and class dependent. Although very few, there are studies that have attempted the subject-independent approach for channel selection [34], [35]. However, experiments in [34] are limited, and [35] is the deep learning model hence the channel selection is dependent on the labels for training the model. Similar is the case with the other deep learning based channel selection models that they are either subject dependent or the task dependent [12], [36]. The critical challenge faced by these channel selection models that beg for a solution is that their selection criteria are dominantly dependent on the specific subject and task at hand. This raises concerns regarding the broader applicability of these models, prompting questions about their ability to generalize effectively to larger and more diverse datasets. We hypothesize that the mutual information between channels can however be strong scaffolds for the subject-independent channel selection. To the best of our knowledge, there has not been any attempt to devise such a subject-independent model.

An additional unsettled issue that got our attention during this study is that several works are showing that a subset of the electrodes of EEG can outperform ‘all channels’ in classifying the stimulus [37]–[39]. On the other hand researchers in [11], [36], empirically show that a subset of the channels is not significantly outperforming the set of all the channels for classifying the MI task. However, the dataset used in [11] has only 22 EEG channels which is smaller than other studies showing it does outperform. In this study, we find that there is no evidence to believe that a subset of channels can outperform all the channels.

In this work, we hypothesize that a subset of channels is good if a *measure of divergence* among them is as small as possible. Furthermore, by minimizing the divergence the selected channels shall be fairly subject and task-independent. To accomplish that, we find the Kullback–Leibler (KL) divergence between a reference channel Cz and all other channels at all time incidents. Afterwards, the expectation of the KL divergence is considered as the rank of the channel. The smaller the rank, the greater the importance of the channels. In addition, we find there is no significant gap in decoding accuracy to believe that a subset of channels can outperform all the channels.

The rest of the paper is organized in three sections. In Section II, methodology has been described. In Section III, we have drawn the results and discussions on the proposed method. Section IV concludes the paper.

## II. Methodology

### A. Notations and Conventions

The general convention is that the bold faced capital letters are matrices and bold small letters are vectors, while italic capital letters are mostly probabilities and random variables with a few exceptions that shall be clear with context. The italic small letters are scalar variables. The indices are represented with small letters that run from 1 to the capital letters. For example, *t* ∈ {1, 2, …, *T*} and *s* ∈ {1, 2, …, *S*}.

### B. Background and Formulation

The basic idea is to rank each channel based on how minimally it is divergent from other channels in the group of selected channels. However, if we find the divergence between every possible channel pair, the noisy channel could be selected. One idea to narrow down the search is to find the divergence of channels from one reference channel and we can fairly assume that if two channels are less divergent from the reference channel, those two channels are themselves less divergent from each other.

The issue is which channel should be considered as a reference channel. A good reference channel would be that which has the least impedance and noise, but, there is little to no literature dedicated to channel-wise noise and impedance analysis. However, in almost all existing channel selection algorithms across all kinds of tasks, Cz has been consistently common as one of the first few selected channels [10], [15], [36]. Another reason for choosing it as a reference is that it is often considered as reference channel in bipolar EEG recordings. Since impedance is measured using this reference channel during the setup, usually people place such electrodes with extreme caution and make sure its contact with the scalp well [40]. Therefore, Cz will be a good choice as the reference channel. Henceforth, a sense of divergence will be evaluated from this reference for each channel and then channels will be ranked based on that. Once the channels are selected, filtering is done followed by classification. Fig. 1 summarises this flow. To formulate the problem of finding the divergence among channels, let us set the notations. Let 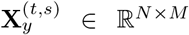 be the EEG signal of *t*^th^ trial of *s*^th^ subject recorded during *y*^th^ stimulus, where *N* is number of channels and *M* is the number of samples of a trial. We can ignore *y* for the channel selection part because the proposed model is independent of it. Hence, it can be described as 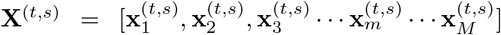, where, 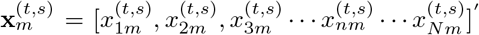, where, 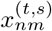 is EEG signal of *n*^th^ channel and *m*^th^ sample. Let the underlined probability space of the EEG signal is (Ω, ℱ, *P*). And 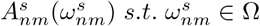 is the random variable representing the EEG signal of *n*^th^ channel and *m*^th^ sample of *s*^th^ subject such that,

**Fig. 1.**
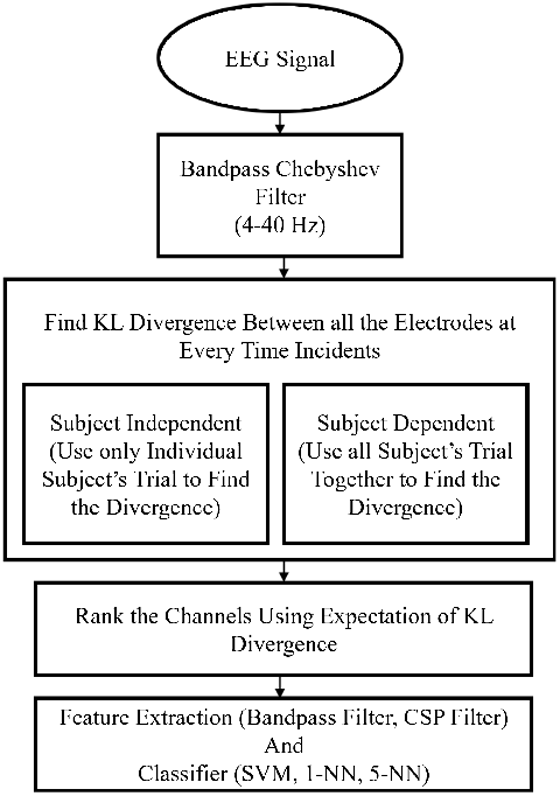
This figure shows the flow of the methodology. The EEG signal will be first filtered using a bandpass filter. Afterward, channel selection is done followed by the classification stage. In channel selection methods we use KL divergence over all the trials to rank the channels. Depending on how this divergence score is evaluated, the proposed method is subject-dependent or independent to it. Finally, we used minimal level of feature reduction using CSP filters, before evaluating our method based on classification accuracy.

**Fig. 2.**
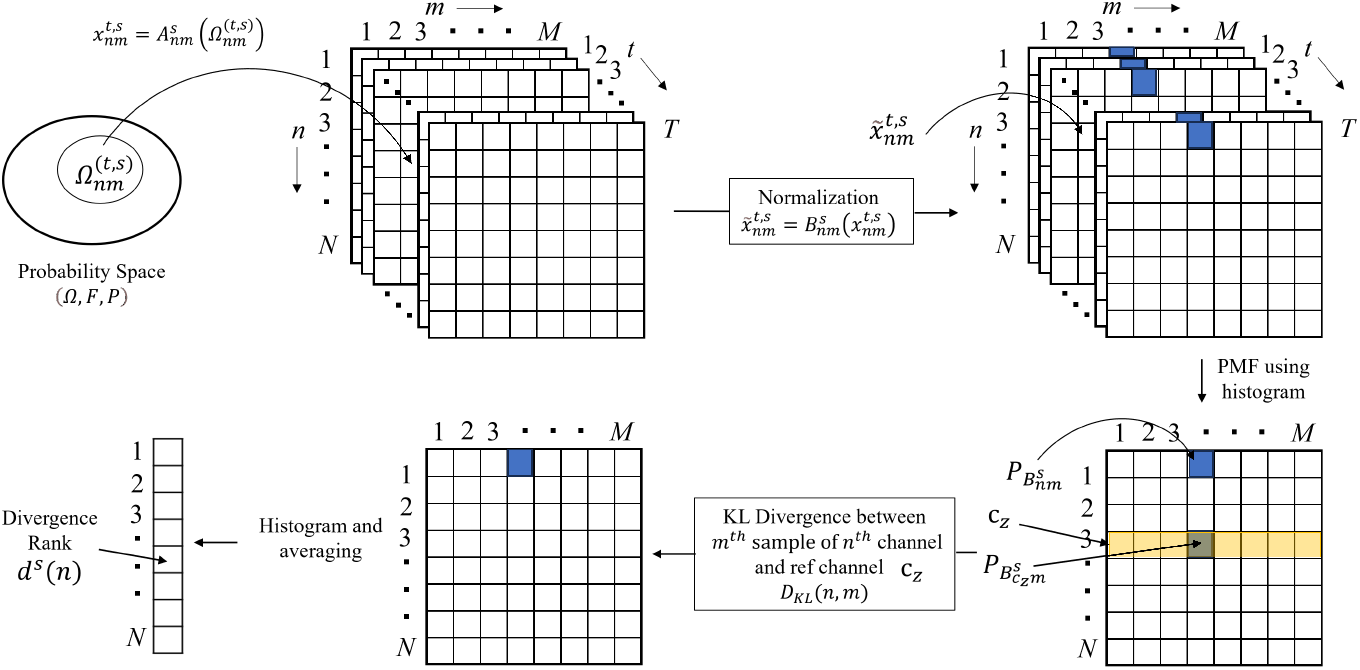
In this figure, the methodology for channel selection has been elaborated pictorially. First, we normalized the data, followed by evaluating the PMFs of each sample and channel using the histogram of all the trail data. Thereafter, we take the KL divergence between a channel and the reference channel and finally average the KL divergence over all the samples to find the divergence rank for the channel.

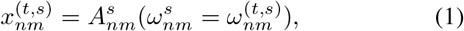

where, 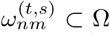 is the probabilistic event that generate the EEG signal at *t*^th^ trial.

EEG signals often contain spurious signals. Such spurious signals can affect the divergence between the channels, significantly. Therefore to attenuate them we used a logarithmic function to normalize the signal. The normalized signal 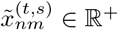 can be evaluated as,

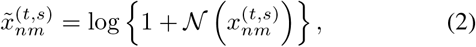

where. *N* (.) is 0 − 1 normalization function along all the samples *M*. The 0 −1 normalization would help us to prevent the saturation of the signal 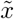 because of the log function.The random variable 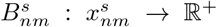 representing the normalized signal shall therefore be defined such that 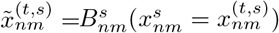.

Next, to find the divergence among the channels we have to estimate the probability mass functions (PMFs) over the random variables 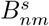 corresponding to the normalized EEG signal 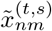 for all possible *n, m* and *s*. Therefore we will be estimating a total of *N* × *M* × *S* number of PMFs. Let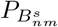 be the PMF of the random variable 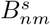. To estimate the PMF 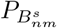 we have *T* number of the trail of EEG signal for each possible *n, m*, and *s*. The number of trials *T* is not very large, hence, a non-parametric model will be more appropriate to estimate the probabilities. It will not be a significant computational burden. Hence, we define PMF 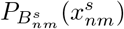 over random variable 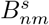using histogram with *H* number of bins. Therefore the half bin size (centers of histogram) *c* = log(2)*/*2*H*, since maximum value 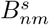can take is log(2). Hence, the PMF 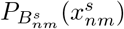 is defined as,

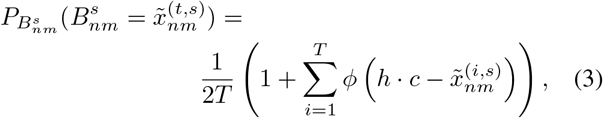

where,

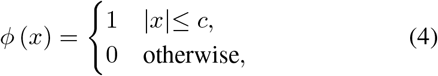

and,

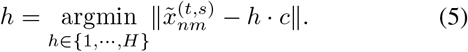

Eqs. 3-5 defines the histogram over the EEG signal 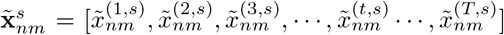 to estimate the PMF 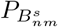at for each channels *n*, samples *m* of each subject *s*. Eq. 5 find the closest bin index of the signal and eq. 3 count the number of EEG trials in that bin. This enables us to estimate all the PMFs. We will use these PMFs to find the divergence between a channel and the reference channel. The divergence can be subject-dependent and independent and hence the channel selection algorithm based on it has been described in the following subsections as subject-dependent and subject-independent.

### C. Subject dependent Channel Selection

Mutual information, divergence, and several other derived measures have been used in literature to rank the EEG channels for channel selection or feature selection [19]–[21]. However, they face limitations as discussed in the first section. In this work, we shall use the expectation of KL divergence between the channel and reference channel to find its rank. Let 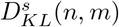 be the KL divergence between the *n*^th^ channel and Cz for the *m*^th^ sample and *s*^th^ subject. It is evaluated as,

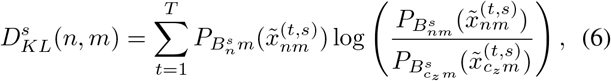

where, 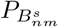is PMF of *n*^th^ channel, *m*^th^ sample and *s*^th^ subject as defined in eq. 3. The ranking of the channel will be done based on an average of the KL divergence over all the *M* samples. To avoid the spurious value of 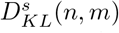affecting the average, we find the histogram and evaluate the sum of the centers weighted with their counts as the divergence metric between the channel and reference channel. Let *d*^*s*^(*n*) be the divergence metric between *n*^th^ channel and the reference channel Cz for the *s*^th^ subject. It is defined as,

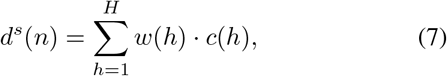

where *w*(*h*) and *c*(*h*) are histogram counts and centers of *M* KL divergences between *n*^th^ channel and Cz. The histogram has been evaluated over samples (time incidents).

Using eqs. 6–7, we can evaluate a divergence score for every channel of every subject. Finally, the channels will be selected based on how small this divergence score *d*^*s*^(*n*) of the channel is. The channel set *C*_*N*_ = {*c*_1_, *c*_2_, *c*_3_, …, *c*_*n*_, …, *c*_*N*_} is sorted such that *d*^*s*^(*n*) ≤ *d*^*s*^(*n* + 1) ∀ *n*. If we need to select best *K* channels then the selected channel set *C*_*K*_ = {*c*_*k*_ |*k*≤ *K*≤ *N*}. The method has been graphically explained in fig 2 for more clarity over the dimensions of the data at every step.

### D. Subject Independent Channel Selection

There are two ways to make the channel selection method subject-independent. One is to take the average over the divergence score *d*^*s*^(*n*) across the subjects (say, average ranks). Let *d*^AR^ is divergence scores of the average rank method, then it can be evaluated as

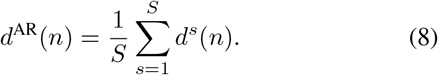

Another way to find the divergence over all the data of all the subjects (say, subject-independent). That can be done by treating all subjects as a single subject and taking the KL divergence over all the trials of all the subjects. Basically, in formulation the number of trials will be *T*× *S* lets call it 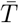 and its index 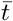. A random variable *Ā* represent the EEG signal such that, 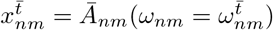, where, 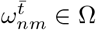, is the event that generate the EEG signal at 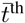 trial. Similarly, let 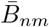 be the random variable representing normalized signal 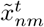. Let PMF over the random variable 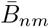 is 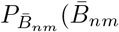 evaluated similarly, except for all the trials 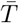, then, subject independent KL divergence *D*_*KL*_(*n, m*) between a channel and reference channel is evaluated as,

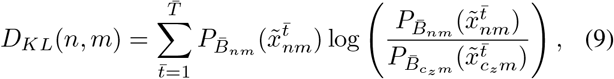

therefore, the divergence rank *d*^INDP^(*n*) for subject independent method for *n*^th^ channel is,

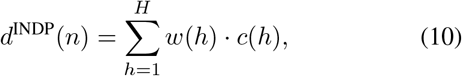

where, *w*(*h*) and *c*(*h*) are histogram counts and centers of KL divergence *D*_*KL*_(*n, m*) between *n*^th^ channel and Cz, similar to as in eq. 7 for the subject dependent version. Finally, we can rank the channels as per the value of *d*^INDP^(*n*) and *d*^AR^(*n*) and select the channels with minimum divergence score.

### E. Inclusion of C3, C4 and Cz

The existing literature shows that 3C (C3, C4, and Cz) channels are the best choice if you want to choose just three channels and are most frequently selected [10], [15], [26], [41]. However, we did not find any convincing argument supporting that except for empirical evidence that suggests strongly as such. We compared 3C with randomly selecting channels and it outperforms all by a great margin (can be seen in fig. 9 & 10). The probable explanation could be lying in the impedance distribution of the channels and scalp, however, we find little to no literature on impedance distribution on electrodes. Nevertheless, 3C is empirically the best 3-channel selection. Thus, we included it as the first three channels for all proposed methods and used the divergence measure for selecting from the 4^*th*^ channel onwards.

### F. Feature Extraction

The goal of the work was to develop a channel selection algorithm and test it using the most basic feature extraction methods for EEG and finally classification. In literature [2], the most common and basic feature extraction pipeline has three stages viz. a) filterbank with overlapping frequency bands, b) common spatial pattern (CSP) Filtering, c) feature Reduction (selection) using mutual information (between features and labels). However, experiments showed no significant effect of using a filterbank over a single filter. With minimal CSP components, feature reduction is unnecessary and relevant only when using a filterbank. Therefore we have used two-stage simple feature extraction stages. The most commonly used frequency band used in MI classification using EEG is 4 −40 Hz [42] therefore we used 10^th^ order Type II Chebyshev filter with cutoff frequencies of 4− 40 Hz. Secondly, we used the CSP filter with 2 components to keep the model overall computationally efficient. For the implementation of the Type II Chebyshev filter and CSP filter stages, we replicated the part of the work presented in [11] with changes for our suitability.

### G. Classifier Pool

We used three classifiers to perform the experiments viz. a) SVM, b) 1-nearest neighbor (1-NN), and 3) 5-NN. The reason to chose 1-NN is that it is proven that it has a theoretical bound on its error, which is twice as much of Bay’s classifier [44]. Therefore, to have a simple check on SVM and 5-NN to be not overfit, we have 1-NN.

The training protocol is that, first, the datasets are divided into 80:20 training-testing sets. Afterward, the training dataset is split into 10 folds. Then SVM, 1-NN, and 5-NN are trained and validated in the 10-fold cross-validation manner for each subject. The best model is selected that has maximum accuracy over the validation set.

## III. Experiments and Results

### A. Dataset Description

We used two publicly available datasets viz BCI competition III, IVa Motor Imagery dataset (BCICIVa) [43], [45] and PhysioNet EEG Motor Moverment/Imagery dataset (PhysioNet) [46], [47]. The former is a rich dataset because it was recorded on 118 electrodes and there are a total of 280 trials per subject. It is useful to judge motor imagery classification models because of its high-density electrodes and a suitable number of trials for training classification models. On the other hand, the latter one is rich because the number of subjects is 109, although electrode density is average (64), hence it will be useful for judging a model that claims to be subject-independent.

1. *BCI competition III dataset IVa:* The dataset [43], [45] is a very rich dataset for channel selection since it has a large number of channels 118. The dataset was recorded at 1000 Hz. The data was recorded for five healthy subjects while visual cues of two classes viz. right hand and right foot (originally for three cues) were presented for 3.5 seconds, with random intervals for a total of 280 trials per subject. For our purposes, we clipped the data at the onset sample till 3500 samples from the onset.
2. *PhysioNet:* In this dataset [46], [47], 64 channel EEG was recorded for 109 subjects performing various MI tasks. Originally, there were 14 runs, out of which we used 3 runs 4^th^, 8^th^, and 12^th^ corresponding to *Task 2 (imagine opening and closing left or right fist)*. The publicly available version of the dataset is at 160 Hz with each trial duration of 4.1 seconds. The number of trials per run is 30 out of which 15 is of MI stimulus and 15 is of resting. We excluded 15 subjects because of the data incompleteness. The excluding criteria are - a) the number of events in any of the three runs is less than 30, b) if any of the trials is recorded duration is less than 4.1s then the whole subject data is excluded, and c) if the number of samples in any run is less than 19200 (160 Hz ×4 seconds ×30 runs) the subject has been discarded. Based on these criteria data of subjects 34, 37, 41, 51, 64, 72, 73, 74, 76, 88, 89, 92, 100, 102, and 104 was discarded. So, we used data of 94 subjects, 3 runs each with 15 trials of 640 samples (4 seconds from onset of the stimulus).

### B. Parameters Tuning and other Details for Replicability

There are several parameters to be tuned. To start with, we decided *H* = 10 to devise the histogram in eq 8. Therefore the size of *w*(*h*) will be *H* × 1. We find the PMF using MATLAB `hist()` with 10 bins. We added 1 to all the counts while finding the PMF so that in case if count in any of the bins is zero, it does not create `NaN` or `inf` in KL divergence since it has the log function. In bandpass filtering, we used 4 − 40 Hz. The order of the filter was 10 since we found it computationally efficient enough for our purposes. For CSP filtering, we used two components of the CSP filter corresponding to the highest and lowest eigenvalues both corresponding to both classes, hence in total 8 components, similar as in [11]. Hence at the input end of the classifiers, the feature dimension is 8.

### C. Empirical Results

Assessment of proposed methods – subject-dependent (Sub Dep), average ranks (Avg Ranks), and subject-independent (Sub Indp) based on MI classification accuracy on both datasets have been drawn in TABLEs I & II and figs. 3, 4, & 6. We compared our results with three existing methods (its subject-independent version) CSP Rank [28], Fishers’s score (FS) [9], [48]–[50] and NMI [20]. The comparison results have been shown in figs. 9 & 10 for both datasets. Comparison of the results has been drawn in detail in Section III-D.5.

**TABLE I.** This table shows the empirical results of the subject-independent methods on the BCICIVA datatset [43]. Results show that the proposed method outperform classical methods CSP Rank [28] and NMI [20] significantly. It performs 15.21% more than 3cs and just 2.91% less than all channels accuracy with as few as 20 channels out of 118. Results in bold are the best among the 3Cs, CSP Rank, NMI, and Sub Indp. All channel accuracy has been provided for reference.

| Subjects | C3, C4 & Cz |  |  | CSP Rank [28] |  |  | NMI [20] |  |  | Sub Indp (20 Channels) |  |  | All Channels |  |  |
| --- | --- | --- | --- | --- | --- | --- | --- | --- | --- | --- | --- | --- | --- | --- | --- |
|  | SVM | 1-NN | 5-NN | SVM | 1-NN | 5-NN | SVM | 1-NN | 5-NN | SVM | 1-NN | 5-NN | SVM | 1-NN | 5-NN |
| aa | 64.53 | 64.14 | 64.14 | 83.85 | 74.89 | 74.89 | **92.04** | 82.07 | 80.68 | 88.64 | **83.46** | **83.46** | 89.64 | 88.25 | 86.86 |
| al | 85.75 | 85.75 | **88.02** | 85.75 | 84.50 | 83.68 | 90.29 | 84.30 | **88.02** | **93.19** | **87.20** | 87.20 | 96.28 | 96.28 | 97.73 |
| av | 66.31 | 51.04 | 56.83 | 66.53 | 59.92 | 63.64 | 68.61 | 64.26 | 66.53 | **76.24** | **71.08** | **67.36** | 77.89 | 70.45 | 68.18 |
| aw | 65.51 | 57.53 | 57.53 | 85.99 | 87.50 | 82.33 | 91.82 | 84.69 | 90.51 | **96.99** | **93.97** | **93.97** | 98.49 | 95.68 | 93.53 |
| ay | 89.59 | 78.70 | 76.67 | 85.53 | 78.22 | 80.09 | 83.50 | 85.37 | 85.37 | **92.69** | **96.42** | **92.69** | 100.00 | 96.58 | 94.71 |
| Average | 74.34 | 67.43 | 68.64 | 81.53 | 77.01 | 76.92 | 85.25 | 80.14 | 82.22 | **89.55** | **86.43** | **84.94** | 92.46 | 89.45 | 88.20 |
| STD | 10.97 | 12.96 | 12.03 | 7.54 | 9.64 | 7.29 | 8.88 | 8.02 | 8.49 | **7.16** | **8.97** | **9.57** | 8.10 | 9.99 | 10.62 |

**TABLE II.** In this table, we show a comparison of empirical results of the subject independent (Sub Indp) PROPOSED METHOD ON THE PhysioNet dataset [46]. Results are compared with a) averaged over 10 times randomly selected channels (RAND), b) CSP rank [28], C) NMI [20], and, d) FS [9] for different numbers of channels. The shown results are averaged over best performing 35, 50, 85, and all (94) subjects. Results in bold are the best SVM classification accuracy among all the methods. The results show that the proposed model outperforms the existing models most of the time.

|  | 35 Subjects |  |  |  | 50 Subjects |  |  |  | 85 Subjects |  |  |  | 94 Subjects |  |  |  |
| --- | --- | --- | --- | --- | --- | --- | --- | --- | --- | --- | --- | --- | --- | --- | --- | --- |
|  | 10 Chs | 25 Chs | 40 Chs | 55 Chs | 10 Chs | 25 Chs | 40 Chs | 55 Chs | 10 Chs | 25 Chs | 40 Chs | 55 Chs | 10 Chs | 25 Chs | 40 Chs | 55 Chs |
| RAND | 75.37 | 87.03 | 92.46 | 95.56 | 72.89 | 85.15 | 90.82 | 94.01 | 68.24 | 80.70 | 86.23 | 89.62 | 66.81 | 79.01 | 84.46 | 87.48 |
| CSP [28] | 83.91 | 93.67 | 98.77 | 99.72 | 79.42 | 90.23 | 95.93 | 97.43 | 70.89 | 83.00 | **89.99** | 91.09 | 66.93 | 80.83 | **87.36** | 87.58 |
| NMI [20] | **83.92** | **94.81** | 97.93 | 99.56 | **80.41** | 91.71 | 95.57 | 97.35 | **71.20** | 83.65 | 89.60 | 91.16 | 67.44 | 81.09 | 86.72 | 88.67 |
| FS [9] | 81.07 | 93.81 | 97.53 | 99.85 | 77.46 | 90.61 | 94.97 | 97.62 | 68.98 | 83.27 | 89.13 | 92.03 | 65.20 | 80.76 | 86.97 | 89.02 |
| Sub Indp | 82.17 | 94.63 | **99.18** | **100.00** | 78.58 | **91.93** | **96.75** | **97.82** | 70.98 | **84.09** | 89.72 | **92.34** | **67.61** | **81.53** | 87.03 | **90.12** |

**Fig. 3.**
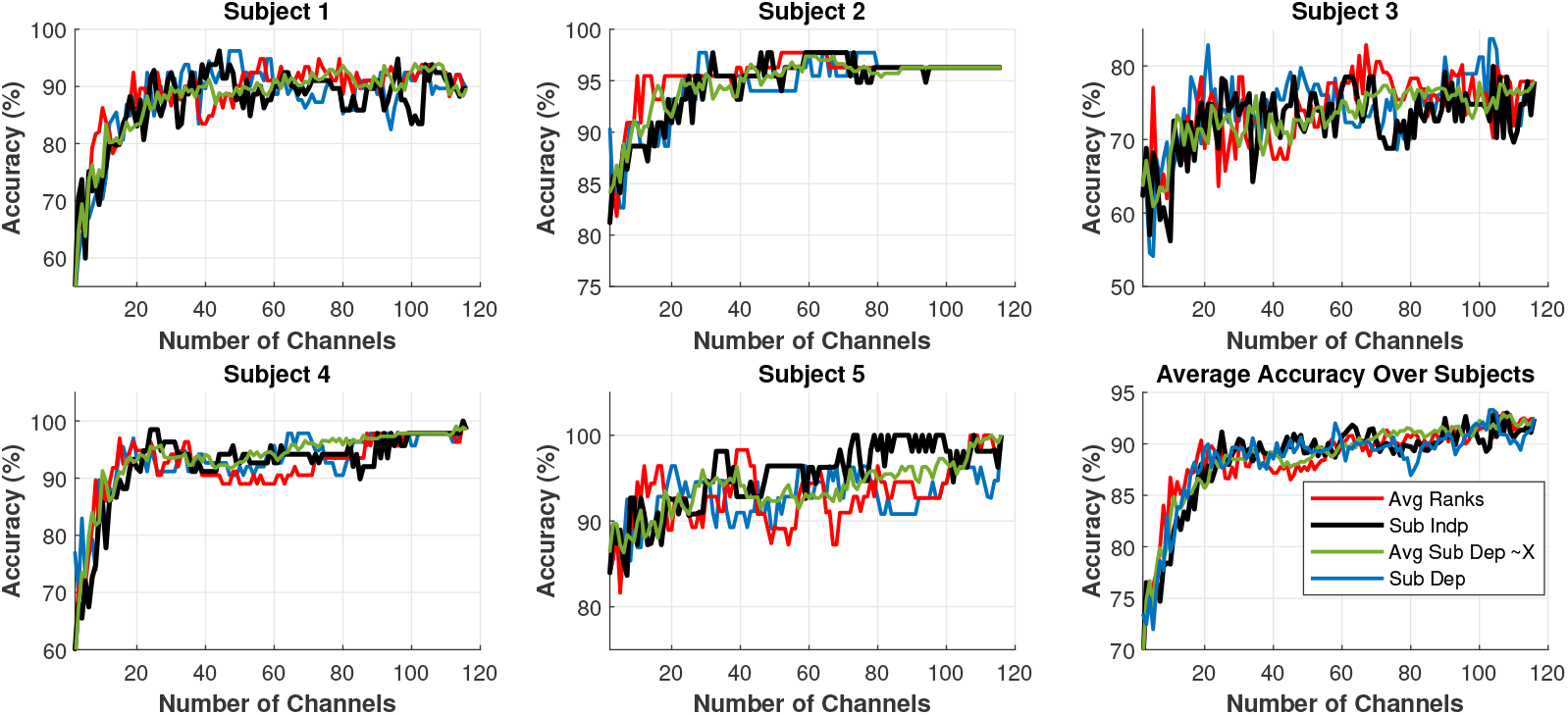
This plot shows how classification accuracy varies with different numbers of channels on the BCICIVa dataset [43] without using bandpass filter stage for channel selection. Four channel selection methods have been compared in this figure viz. a) the subject dependent (Sub Dep) method where the ranks for the channels have been evaluated using the EEG signals of just that subject, b) average rank (Avg Rank) is just the average of the individual subject’s ranks, c) subject independent (Sub Indp) method, we used all the trials together of all the subjects to find the ranks of the channels, and d) Avg Sub Dep X is average of the performance of channel selection based on other subject EEG data. It can be seen that *Avg Rank* and *Sub Indp* perform better than subject-dependent models most of the time with the smaller number of channels, however, subject-dependent methods perform better with **80 *−* 100** channels.

**Fig. 4.**
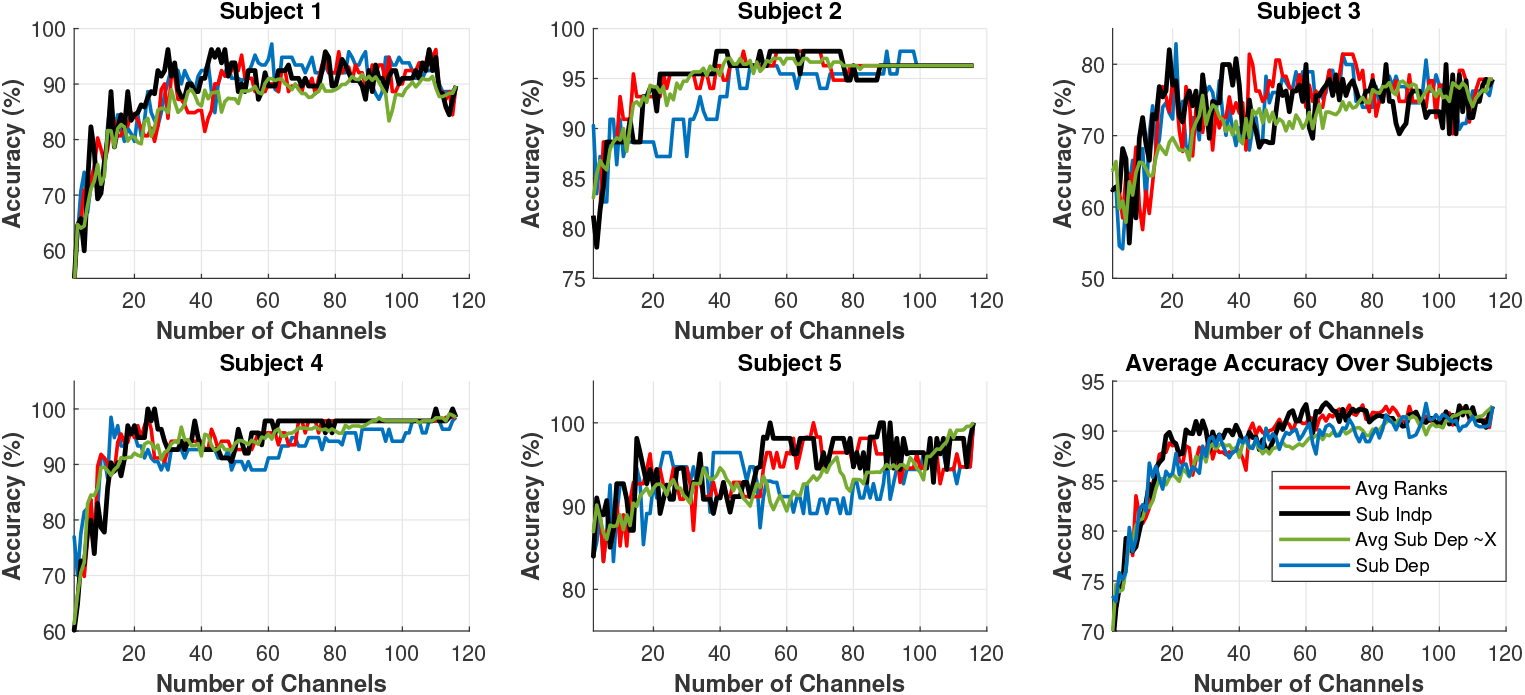
In this plot, how classification accuracy varies with different numbers of channels on the BCICIVa dataset [43] using bandpass filter stage for channel selection have been shown. The most important point to notice in this plot is that the accuracy plot is more stable and consistent than that of *without filters* in fig. 3. Most clearly it can be seen with plots of *Sub Indp* plots of Subject 2 & 3. However, overall with or without filters, *Sub Indp* method performs the best among all for the smaller to average number of channels.

TABLEs I & II shows that the proposed subject-independent method outperforms the existing models. We will be looking at the performance based only on the SVM classifier, for discussion purposes. Classifiers 1-NN and 5-NN have been used to see if kernel SVM does not behave abnormally. It can be seen that from the TABLE I, on the BCICIVa dataset, as few as 20 channels (approx. 17% of all channels) perform almost 15.21% more than 3C and just 2.91% less than *all channels*. Similarly, for the PhysioNet dataset, the proposed subject-independent outperforms the existing channel selection models as can be seen in the TABLE II. It shows that for different numbers of best-performing subjects, the proposed model consistently outperforms, indicating its generalizability over subjects. In addition, the standard deviation (in the interest of brevity, it has been provided in supplementary materials) of these accuracies is also lower which further enforces the generalizability of the proposed model.

How the classification accuracy varies with various numbers of channels of all the proposed methods has been shown in figs. 3, 4, & 6. In figs 3 and 4, we have compared four type of classification performance in each plot for the BCICIVa dataset, viz. a) Subject Dependent (Sub Dep) derived from the equation 7; b) Average Rank (Avg Rank) derived from equation 8; c) Subject Independent (Sub Indp) derived from equation 9; and d) Average performance of Dependent on other Subject (Avg Sub Dep X). The first three are straightforward, and the fourth ‘Avg Sub Dep X’ is basically can be explained as follow – let’s say for subject 1 we are evaluating the performance, then we find the ranks of channels using EEG signals of one of the other subjects than 1 and evaluate the performance. Finally take the average over all the rest of the subject. Fig. 3 shows how channel selection worked in the absence of bandpass filtering in the channel selection stage. The observation reveals that channel selection is effective even without filters; however, the method demonstrates increased robustness when filters are incorporated, as can be seen in figs. 4 & 5. Figs. 4 & 6 shows that the subject-independent method performs most consistently for most of the subjects. Another important conclusion that can be drawn from the results in figs. 4, & 6 is that the classification accuracy almost monotonically increases with increasing number of channels. This is in contrast with the studies in several state-of-the-art works on this topic that a set of fewer channels can outperform all the channels in classification [15].

### D. Discussion

In this section, we have discussed how the various aspects of channel selection affect the proposed methods, how the proposed model is subject-independent, and compared the results with the existing models.

1. *Effect of Bandpass Filter on Channel Selection:* We analyzed how channel selection is affected by whether we use signal filtering or not. How much KL divergence based ranking of the channel affected by the filtering? In most EEG processing, the bandpass filter(s) are used which is generally computationally costly, to suppress the artifacts and power line interference. However, we find that the classification accuracy is not heavily affected because of filtering. Fig. 5, shows that difference between accuracy because of filtering is marginal however crucial. Most of the time accuracy is marginally better than that of without filters, however, an important use we can see from the plot is that, with filters, the accuracy curve seems more consistent and hence seems generalizable.
2. *Subject Independence Analysis From Selected Channel Point of View:* The main goal of the study was to devise a truly subject-independent channel selection method. We ranked the channels based on a divergence measure. We analyzed how much the subject-independent channel selection method shares the selected channels with the subject-dependent version of it. Figs. 7 & 8 graphically shows that the subject-independent methods share most of the channels with subject-dependent methods. Fig. 7 refers to the dataset BCICIVa [43], where we can see that, at its peak the number of different channels on average is approximately 15% for the subject independent method. Furthermore, this difference is almost half of it with the average rank method (black plot). A similar inference can be made from the fig. 8 for the PhisioNet dataset [46], where the subject-independent method is different from other methods maximum up to 15% averaged over 94 subjects. It shows that the proposed subject-independent method is generalizable up to a great extent.
3. *Subject Independence Analysis From Accuracy Point of View:* Inferences about how the subject-independent method is independent of subjects can be made from the figs. 4 & 6 for the datasets BCICIVa [43] and PhisioNet [46] respectively. In the first dataset since the number of subjects is just 5 it is easy to visualize that for all the subjects, *Sub Indp* method performs as good as subject-dependent methods for most cases and better than them at the smaller number of channels. Fig. 6 shows that for the smaller number of channels *Sub Indp* method performs adequately well, and performs better for 40 − 64 channels. However, there is an important explanation for this. For up to 30 channels, even classical models do not perform better than randomly selected channels. This may be due to the PhysioNet dataset having only 64 channels, which is less dense compared to the BCICIVa dataset. As a result, the shunting effect might be smaller, and randomly selected channels could perform well because there are more good channels available. Overall we can infer that the proposed method performs adequately without sacrificing on classification accuracy.
4. *Relevancy of Channels and Divergence:* In fig. 4 for the 2^nd^ and 4^th^ subject, we can see that the *Sub Indp* has streigh line while channels varies between 80 −118. This shows that divergence information has been captured by already selected channels and no new addition of enough divergence is brought up by the incoming channel to affect the classification accuracy. This also shows that divergence is an excellent measure for channel selection.
5. *Comparison of Methodology:* The novelty of the proposed model lies in its subject-independent and generalizable nature. Most of the recent models, for one, are subject and task-dependent, and secondly, it is almost impossible to have a subject-independent version of it [15], [36], [41] and yet be fair while comparing with their work. We can classify the channel selection methods into two classes, a) where the channel selection algorithm and classification stage are modular and separate from each other [9], [15], [28], and, b) where both models are entangled and depend on each other [36], [41]. By the nature of the method itself, it is not possible to compare with the latter class. There is another concern about the comparison study in the existing works that those are not compared with randomly selected channels. It should be noted that the comparison of randomly selected channels is not equivalent to the comparison of random classification.We have, therefore, compared our work with classical methods that are steadily extendable to its subject-independent version. We compared our work with three channel selection models FS [9], CSP Rank [28], and NMI [20] in addition to comparing it with the average of randomly selected channels 10 times (Rand Avg).The comparison results are shown in fig. 9 & 10. Fig. 9 shows how the performance of the proposed model *Sub Indp* varies with the number of channels in comparison with the classical models like FS, CSP Rank, and NMI, along with Rand Avg on the BCICIVa dataset [43]. It can be seen that the proposed method has outperformed all the methods significantly. However, this is not so apparent in the case of the PhysioNet dataset as it can be seen in fig. 10. For the 35 −50 number of channels, the proposed model outperforms significantly over all other methods. However, for the smaller number of channels (3 −35) NMI and FS dominate the proposed method. However, for channels 3 −35 the accuracy is steeply increasing and yet to be saturated, hence it may not be the best point of comparison. It can be said that it is ‘yet to be the saturated area’ for classification accuracy for all the methods. To support this reasoning, it can be seen that even with BCICIVa dataset in fig. 9 the proposed method is not a winner in the ‘yet to be saturated area’ (in this case it is much smaller between 3 −15). Overall, the proposed method performs better than the existing methods.
6. *Underlying Assumption on Performance:* As we discussed in the first section, EEG channel selection is an NP-hard problem and therefore at the heart of the problem, some assumptions help narrow down the search. In this work, following are the two major underlying assumptions we have made. The first and foremost important assumption we have made is that irrespective of the spatial location of the electrodes, it captures the signal uniformly which stems from Helmholtz theorem [16]. This is the reason, we have not used montage locations or any neighborhood functional in ranking the channels, in contrast with several recent works [15]. The second major assumption is that the channel Cz is the reference channel. We have made attempts to justify this assumption at length in Section II-B, and discussed possibilities beyond it in Section III-D.7.
7. *Limitations and Future Work:* The proposed work performed well on the two publicly available datasets. It shows a subject independent character up to a great extent. However, there are several fronts where there is scope for improvement. First, based on the classification accuracy point of view, the proposed method did not perform better than the comparing models for the smaller number of channels, and there is scope to improve upon it. Secondly, the proposed approach mainly assumes that the reference channel Cz has low impedance and is a good channel. The reason to select Cz as a reference has been backed by literature, however, if this is not the case, then the proposed method may not perform well. However, there could be a quick fix to use ‘an average of scalp area’ as a reference and the overall principle of using KL divergence can be used intact. Multiple references also can be used for electrodes from different areas of the scalp to capture the potential distribution of the microstates. Another issue is the question of the generalizability of the proposed model over other tasks than MI. Although we have not used the task labels for the channel selection as such in our model, it needs scrutiny and probable modification in the principle for generalizing over tasks. One way to tackle this is to devise a divergence measure between electrodes that is independent of the EEG signal but captures the essence of the task-related topology of the scalp electrodes and thereafter combine it with the KL divergence model.

**Fig. 5.**
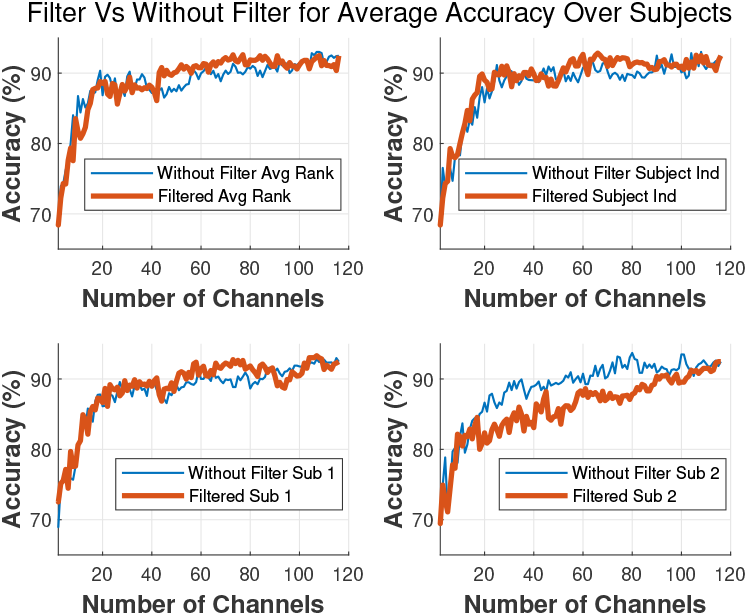
The impact of bandpass filtering on the selection of channels on the assessment of classification accuracy on the BCICIVa dataset [43]. The plots reveal that filtering has a discernible impact on classification accuracy, influencing a smoother and more consistent change in accuracy with the number of channels compared to cases without filters. However, the difference between the accuracy is not visibly significant. There are up to **20** channel differences in the selected channels contributing to variations in the accuracy.

**Fig. 6.**
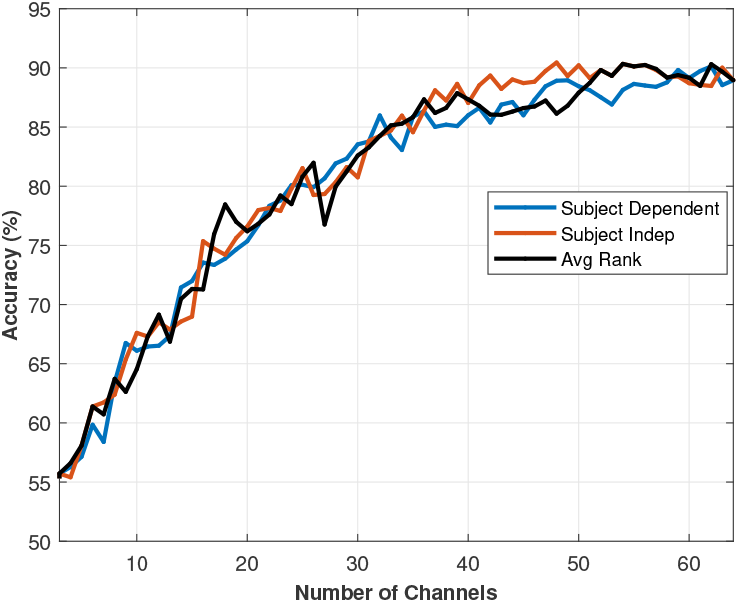
This graph illustrates how the accuracy of classification changes with varying numbers of channels on the PhysioNet dataset [46]. The classification accuracy is averaged over **94** subjects. The shown three methods are subject-dependent, average rank, and subject-independent. Again, it can be seen that on average subject-independent methods perform better (at least as good as) than subject-dependent methods, with the additional advantage of it is more generalizable.

**Fig. 7.**
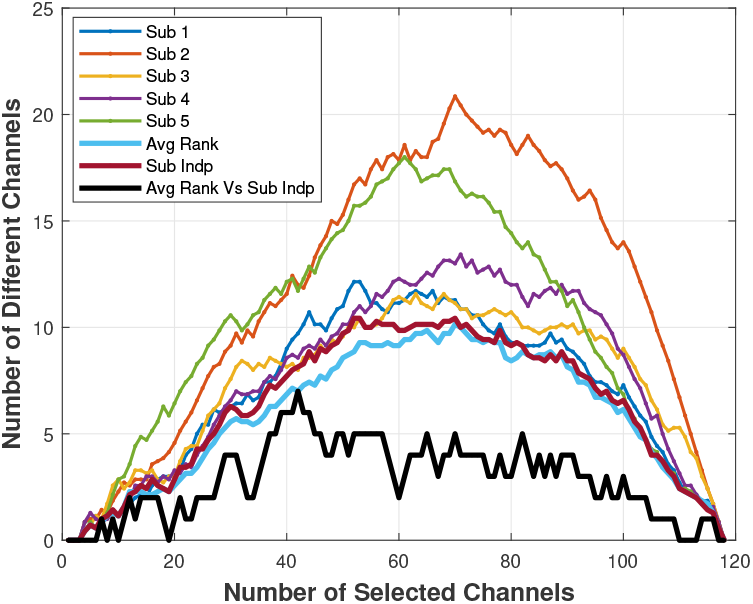
In this plot, we examined the divergence among various proposed channel selection models based on the number of distinct channels selected on the BCICIVa dataset [43]. The associated plot for the method illustrates the average disparity in selected channels compared to all other methods. For instance, *Sub 1* represents the average difference from other methods such as *Sub 2, 3, 4*,… based on the selected channels. It shows that both subject-independent methods are close to each other up to a great extent.

**Fig. 8.**
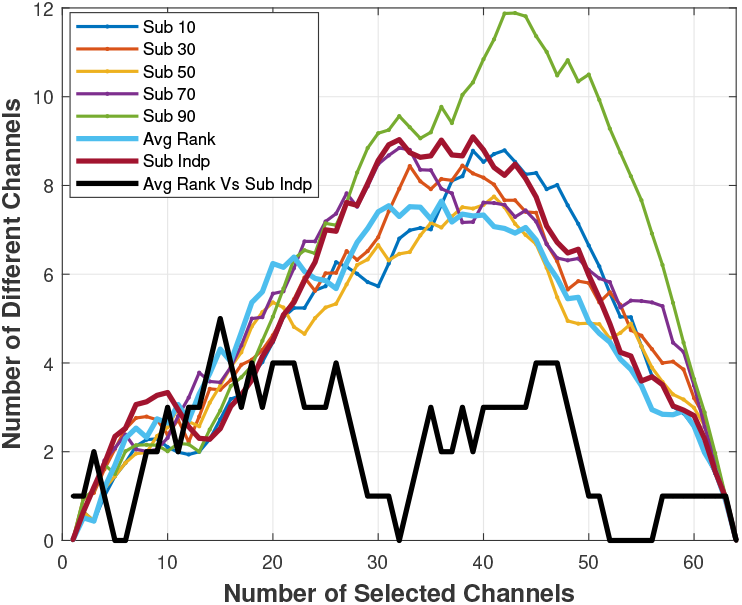
Plot on how all methods are different from each number based on the difference in selected channels for the classification of MI on the PhysioNet dataset [46]. It shows the average of the number of different channels of the method from all other methods. The average is over **96** ranking models (**94** subjects, Avg Rank, and Sub Indp). Again, it can be seen that the subject-independent method is almost similar to the other models and has a peak average difference of less than **15**% of channels, hence it is generalizable up to a great extent.

**Fig. 9.**
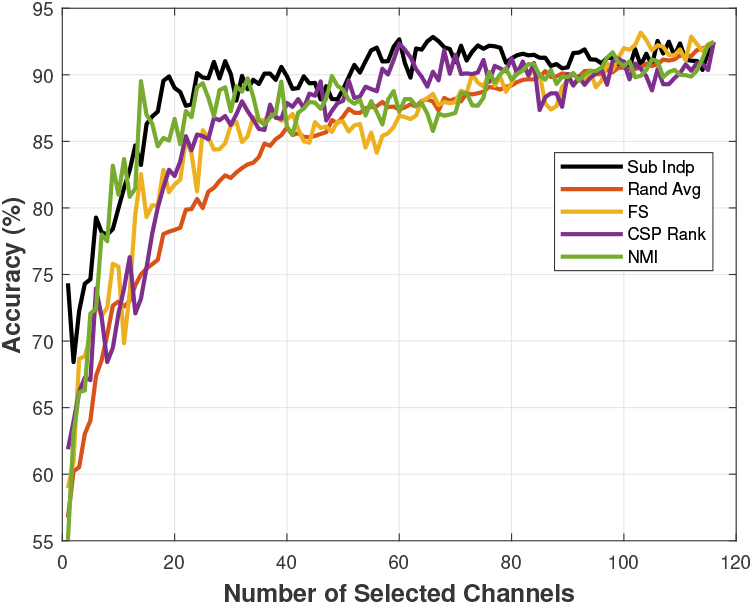
[AVG OVER SUBJECTS] This figure illustrates the comparison between the proposed method and classical models on the BCICIVa dataset [43]. The proposed model is evaluated against three classical models (CSP Rank [28], FS [9], and NMI [20]) as well as randomly selected channels (Rand Avg). The plots distinctly demonstrate the superior performance of the proposed method over all other approaches. Another notable point is that the accuracy monotonically (approximately) increases with the number of channels.

**Fig. 10.**
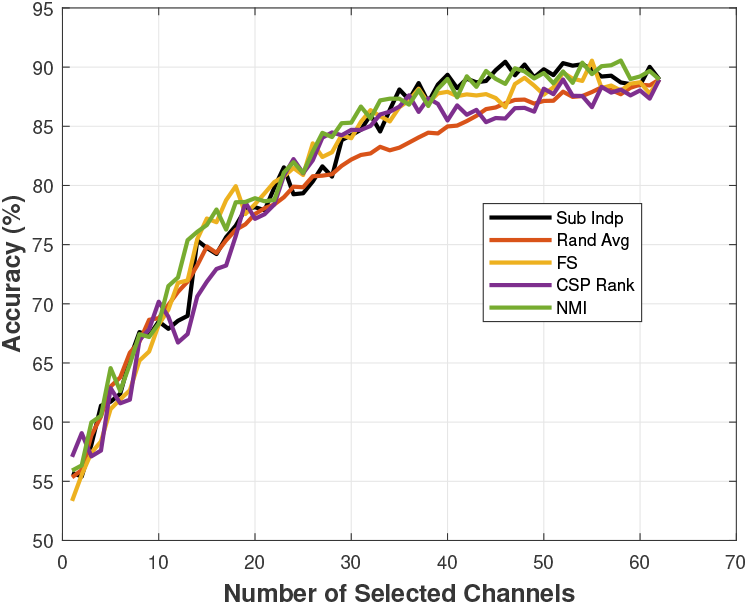
[AVG OVER SUBJECTS] Plot illustrating the comparison of the proposed method with existing models in terms of classification accuracy on the PhisioNet dataset [46]. The results have been compared with three classical models viz. CSP Rank [28], FS [9], and NMI [20] and randomly selected channels. The results show that the proposed method outperforms CSP Rank, FS, and Rand Avg methods significantly. However, the proposed method underperforms the NMI for channels less than **35** and significantly outperforms afterward.

## IV. Conclusions

In this work, we presented a subject-independent EEG channel selection algorithm and analyzed various aspects of it. We hypothesized that a good way to pursue that would be to use a metric that has a sense of divergence to find a rank for all the channels. We used KL divergence of trials between channels and a reference channel and further their expectation as rank to the channels. We proposed the subject-dependent and independent versions of it and analyzed the differences. We validated our method on two publicly available datasets. We find that the set of channels with smaller divergence among them is a better choice for the motor imagery classification. Experimental results show that the proposed approach performs better than existing models and is generalizable. Future work includes extending the experiments on other tasks than motor imagery to test how it works with other tasks.

## Acknowledgement

We thank Dr. Pasquale Arpaia and Dr. Antonio Esposito for their help in replicating their work, especially with CSP implementation. We also thank Mr. Rohit Misra (MSR, IIT Delhi) for the discussion and debate during the development of our model,

## Data availability

The datasets used in this work are publicly available. The codes used in this manuscript are available at https://github.com/rdevm23/eeg_ch_sel_kl_div/tree/ public for reproducibility.

